# A new cancer-testis long noncoding RNA, the OTP-AS1 RNA

**DOI:** 10.1101/350793

**Authors:** Iuliia K. Karnaukhova, Dmitrii E. Polev, Larisa L. Krukovskaya, Alexey E. Masharsky, Olga V. Nazarenko, Andrei A. Makashov, Andrei P. Kozlov

## Abstract

Orthopedia homeobox (*OTP*) gene encodes a homeodomain-containing transcription factor involved in brain development. *OTP* is mapped to human chromosome 5q14.1. Earlier we described transcription in the second intron of this gene in wide variety of tumors, but among normal tissues only in testis. In GeneBank these transcripts are presented by several 300-400 nucleotides long AI267901-like ESTs.

We assumed that AI267901-like ESTs belong to longer transcript(s). We used the Rapid Amplification of cDNA Ends (RACE) approach and other methods to find the full-length transcript. The found transcript was 2436 nucleotides long polyadenylated sequence in antisense to *OTP* gene. The corresponding gene consisted of two exons separated by an intron of 2961 bp long. The first exon was found to be 91 bp long and located in the third exon of *OTP* gene. The second exon was 2345bp long and located in the second intron of *OTP* gene.

The search of possible open reading frames (ORFs) showed the lack of significant ORFs. We have shown the expression of new gene in many human tumors and only in one sampled normal testis. The data suggest that we discovered a new antisense cancer-testis sequence *OTP*-*AS1* (*OTP*- antisense RNA 1), which belongs to long noncoding RNAs (lncRNAs). According to our findings we assume that *OTP-AS1* and *OTP* genes may be the CT-coding gene/CT-ncRNA pair involved in regulatory interactions.

**Author summary:** Previously, long non-coding RNAs (lncRNAs) were considered as genetic “noise”. However, it was later shown that only 2% of genomic transcripts have a protein-coding ability. Non-coding RNA is divided into short non-coding RNAs (20-200 nucleotides) and long noncoding RNAs (200-100,000 nucleotides). Genes encoding lncRNA often overlap or are adjacent to protein-coding genes, and localization of this kind is beneficial in order to regulate the transcription of neighboring genes. Studies have shown that of lncRNAs play many roles in the regulation of gene expression. New evidence indicates that dysfunctions of lncRNAs are associated with human diseases and cancer.

In our study we found a new cancer-testis long noncoding RNA (*OTP-AS1*), which is an antisense of protein-coding cancer-testis gene (*OTP*). Thus, *OTP-AS1* and *OTP* genes may be the CT-coding gene/CT-ncRNA pair involved in regulatory interactions. This is supported by the similar profile of their expression. *OTP-AS1* may be of interest as a potential diagnostic marker of cancer or a potential target for cancer therapy.

Part of *OTP-AS1* gene (5’-end of the second exon) is evolutionary younger than the rest of gene sequence and is less conservative. This links *OTP-AS1* gene with so-called TSEEN (tumor-specifically expressed, evolutionary novel) genes described by the authors in previous papers.

## Introduction

Earlier we performed *in silico* analysis of the UniGene transcribed sequences database, which includes human transcribed sequences and found that AI267901 sequence is expressed tumor-specifically [1]. Subsequently we confirmed tumor-specific expression of AI267901 experimentally [2, 3].

We mapped the sequence AI267901 to human genome using UCSC Genome Browser. The studied sequence was found to be located on 5q14.1 in the second intron of the human *Orthopedia homeobox* (*OTP*) gene [3].

We suggested that AI267901 and other similar ESTs are the parts of a longer RNA. In order to verify our hypothesis and to obtain complete nucleotide sequence of putative long RNA we used Rapid Amplification of cDNA Ends (RACE) approach.

## Results

We assumed that AI267901 and similar ESTs belong to one long transcript so we used the Rapid Amplification of cDNA Ends (RACE) to find its and 3’ ends. Results of the two-round amplification of the 5’ end of the transcript are presented at Fig 1. The figure shows the PCR-product of 443 bp long. The obtained fragment was further cloned, propagated in *E.coli* and sequenced (S1 Sequence). The resulting sequence was aligned with the human genome. We mapped the 443 bp fragment to the chromosome 5 and found that it consists of two exons (91 and 352 bp) separated by 2961 bp long intron (Figs 2 and 4).

**Fig 1.**
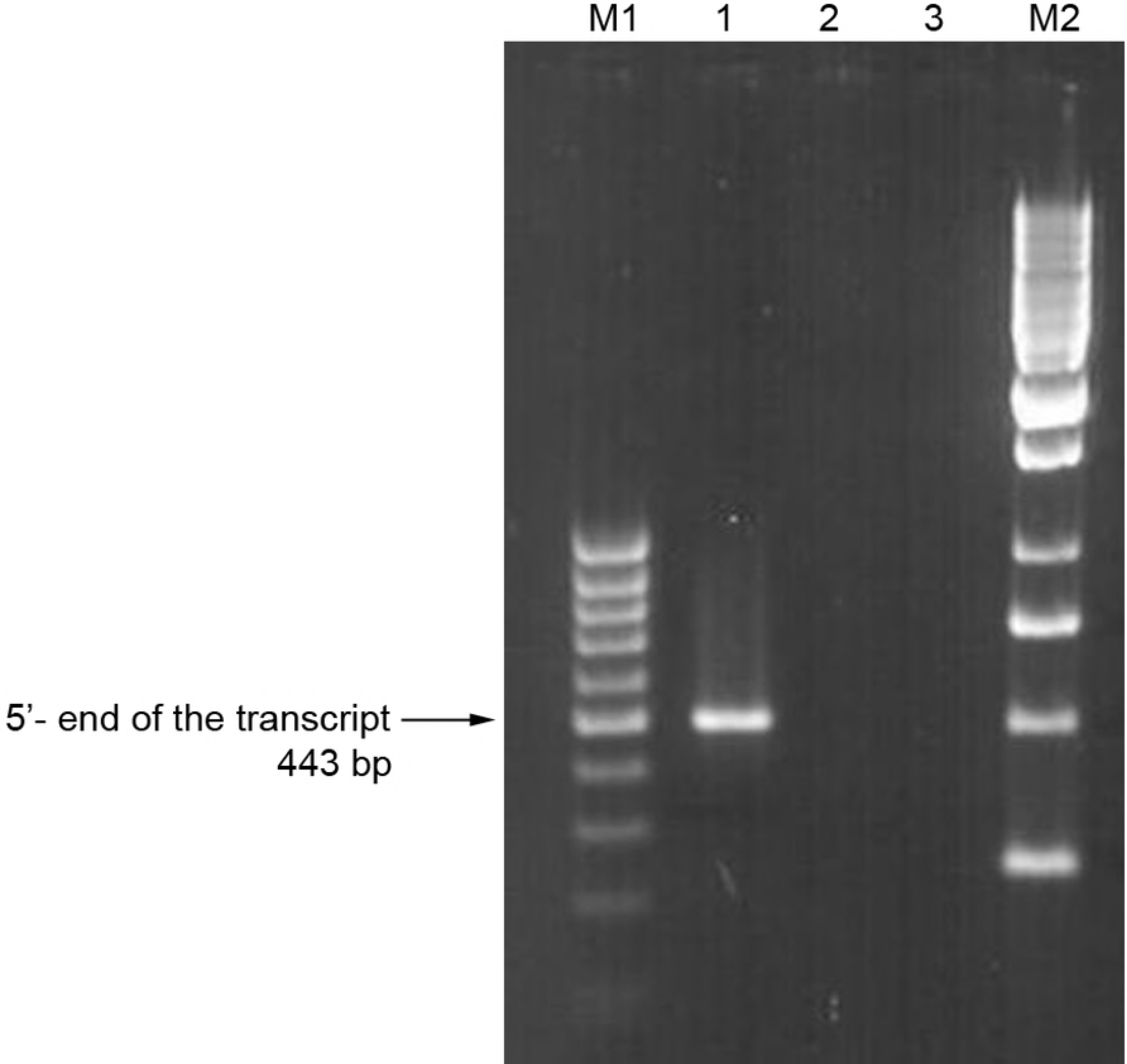
Two-round amplification of the studied gene 5’ end using gene specific and adaptor primers. 1 - adaptor primer and rev(N) 2 - negative control, first round of PCR, no template added 3 - negative control, second round of PCR, no template added M1 - GeneRuler™ 100 bp DNA ladder (Fermentas) M2 - GeneRuler™ 1 kb DNA ladder (Fermentas)

**Fig 2.**
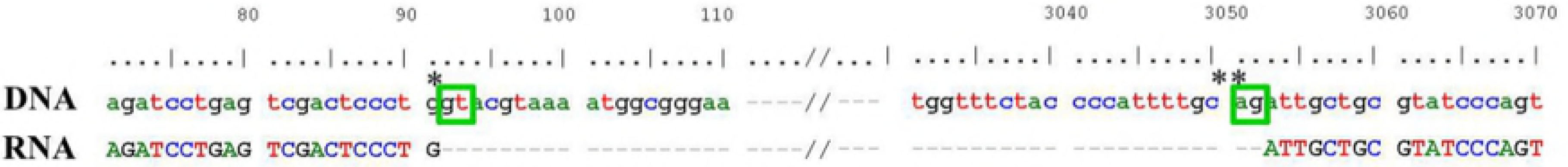
Exon/intron borders of the studied gene. * gt - donor splicing site **ag - acceptor splicing site

Analysis of the exon/intron borders (Fig 2) demonstrated that the studied sequence is encoded by “plus” DNA strand of the chromosome 5, *i. e.* located in antisense to the *OTP* gene. The first exon is located in the third exon of *OTP* gene, and the second one - in the second intron of *OTP* gene.

We were unable to extend the 3’ end of the transcript using the RACE technique. Therefore to determine the 3’ end of the transcript we performed the first strand cDNA synthesis with RT and oligo(dT) adapter primer followed by 2-round PCR. The results are presented at Fig 3. The obtained three fragments of different size were further cloned, propagated in E.coli and sequenced. We found that fragment №1 (see sequence in S2 Sequence) was 2311 bp long, polyadenylated, and located at the 5th human chromosome as an extension of the 443bp fragment found earlier with 318 bp overlapping of the AI267901 sequence (Fig 4). Fragment №2 was found to reside on the 3rd chromosome (chr3:14247460-14247871, data not shown). And, finally, fragment №3 (see sequence in S3 Sequence) was found to be polyadenylated, 940 bp long and also located on the 5th chromosome, fully matching the 3’ end of the fragment №l (Fig 4).

**Fig 3.**
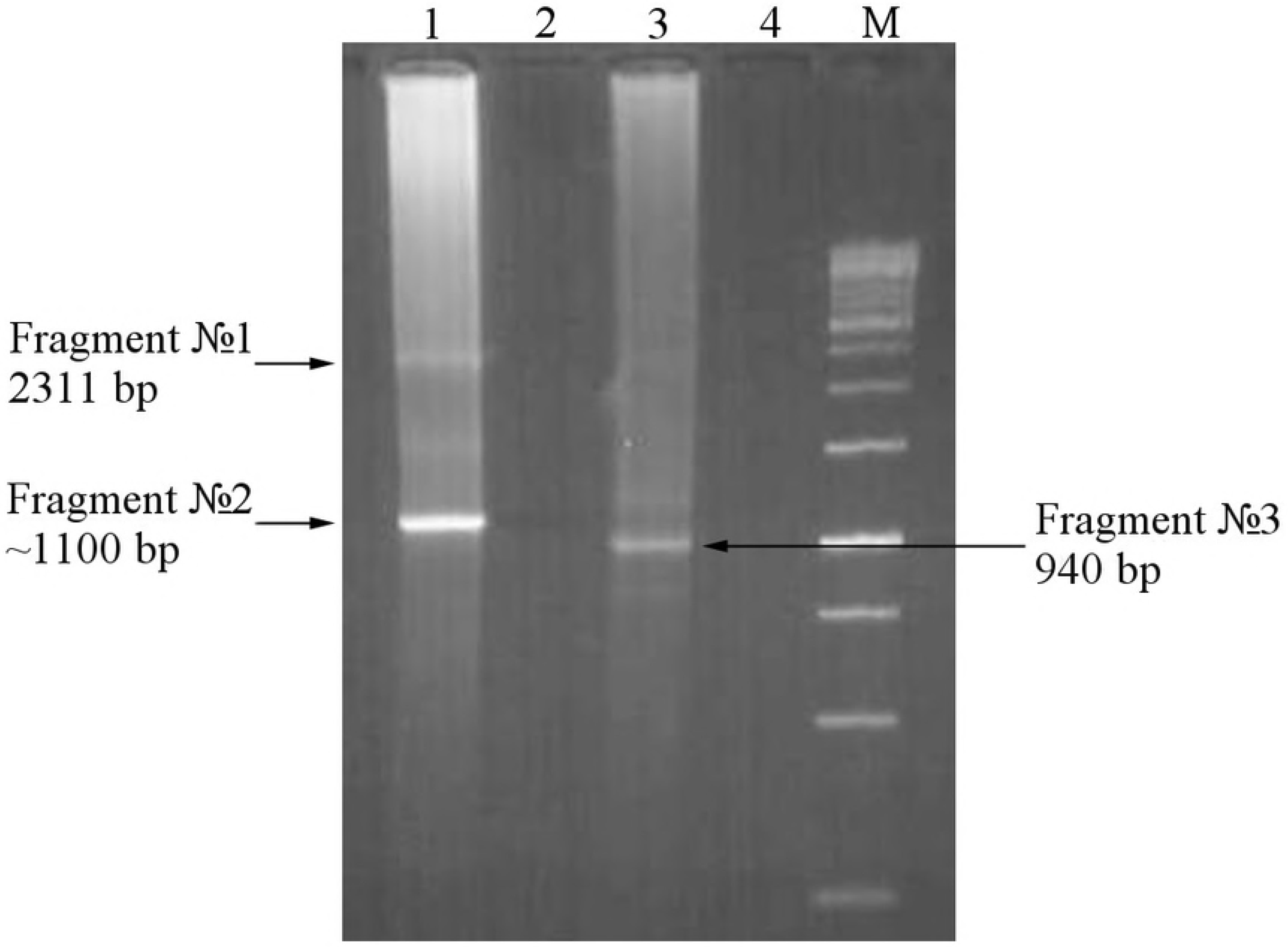
Two-round amplification of the studied gene 3’ end using gene specific and adaptor primers. 1 - adaptor primer and forv1(N) 2 - negative control, first round of PCR, no template added 3 - adaptor primer and forv2(N) 4 - negative control, second round of PCR, no template added M - GeneRuler™ 1 kb DNA ladder (Fermentas)

**Fig 4.**
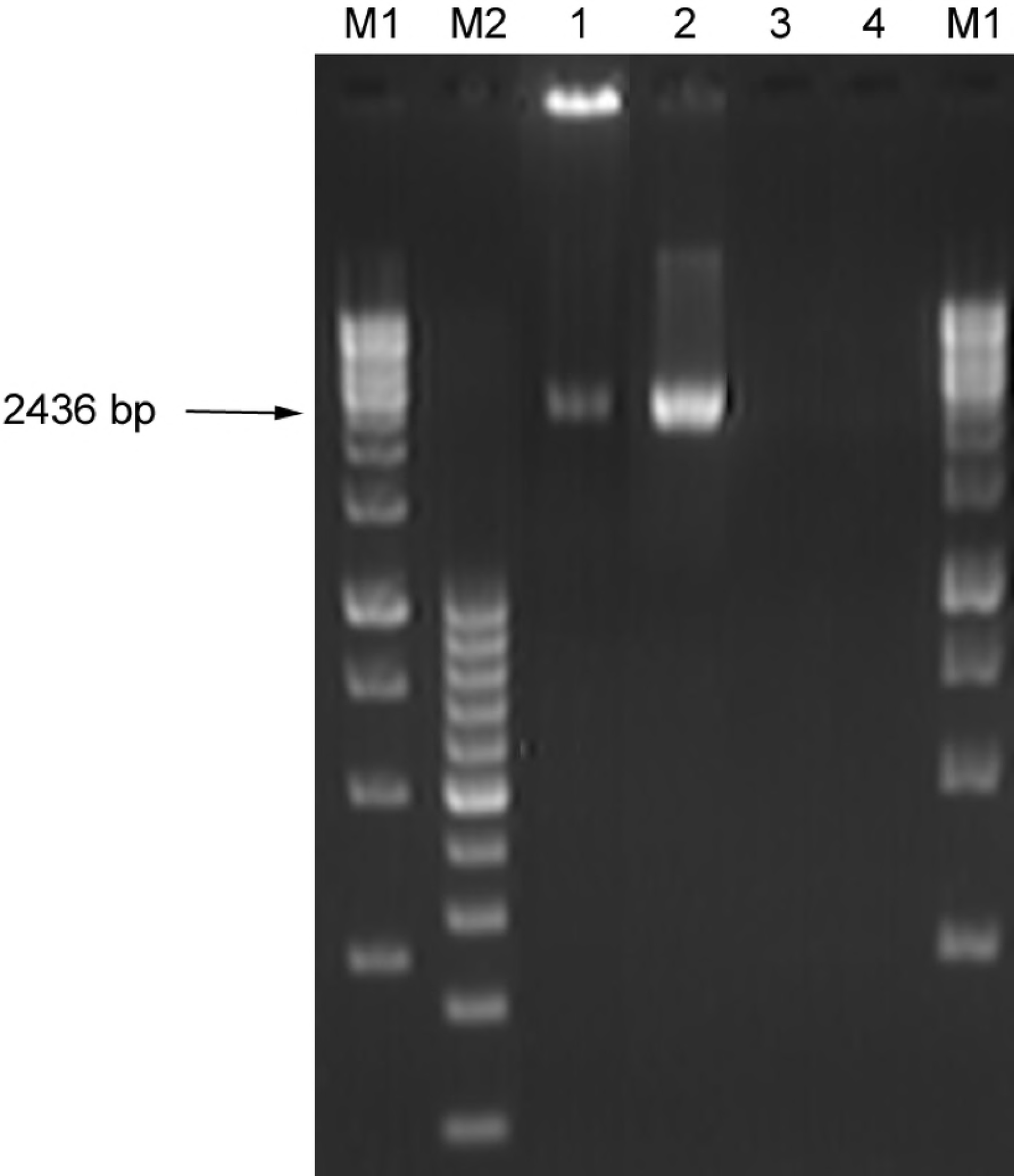
Genome localization of *OTP* gene and its sequenced fragments. Schematic alignment of the sequenced fragments (5’ end fragment - yellow, 3’ end fragment №l - green, 3’end fragment №3 - cyan, assembled full-length sequence - red) with the 5th chromosome (black) and the OTP gene (blue).

Using the BioEdit (v.7.2.5) software and UCSC Genome Browser we determined the sequence of the previously unknown gene located on chromosome 5 in antisense to the *OTP* gene. The corresponding transcript of the gene is 2436 bp long and polyadenylated. This gene consists of two exons: the first exon is 91 bp long and locates in the third exon of *OTP* gene, and the second one is 2345 bp long and locates in the second intron of *OTP* gene. The exons are separated by an intron of 2961 bp long (Fig 4).

To obtain the full length sequence of this gene experimentally the 2-round RT PCR was performed. RNAs isolated from 293T cells or uterus endothelium adenocarcinoma were used as templates. RT was performed with the oligo(dT) primer, PCR - with the full-forv and full-rev primers. The resulting fragment is 2436 bp long (Fig 5) as it was predicted by *in silico* analysis (Fig 4). This full lenght transcript was cloned into TA cloning vector, propagated in *E. coli* and sequenced. Its sequence is presented in S4 Sequence.

**Fig 5.**
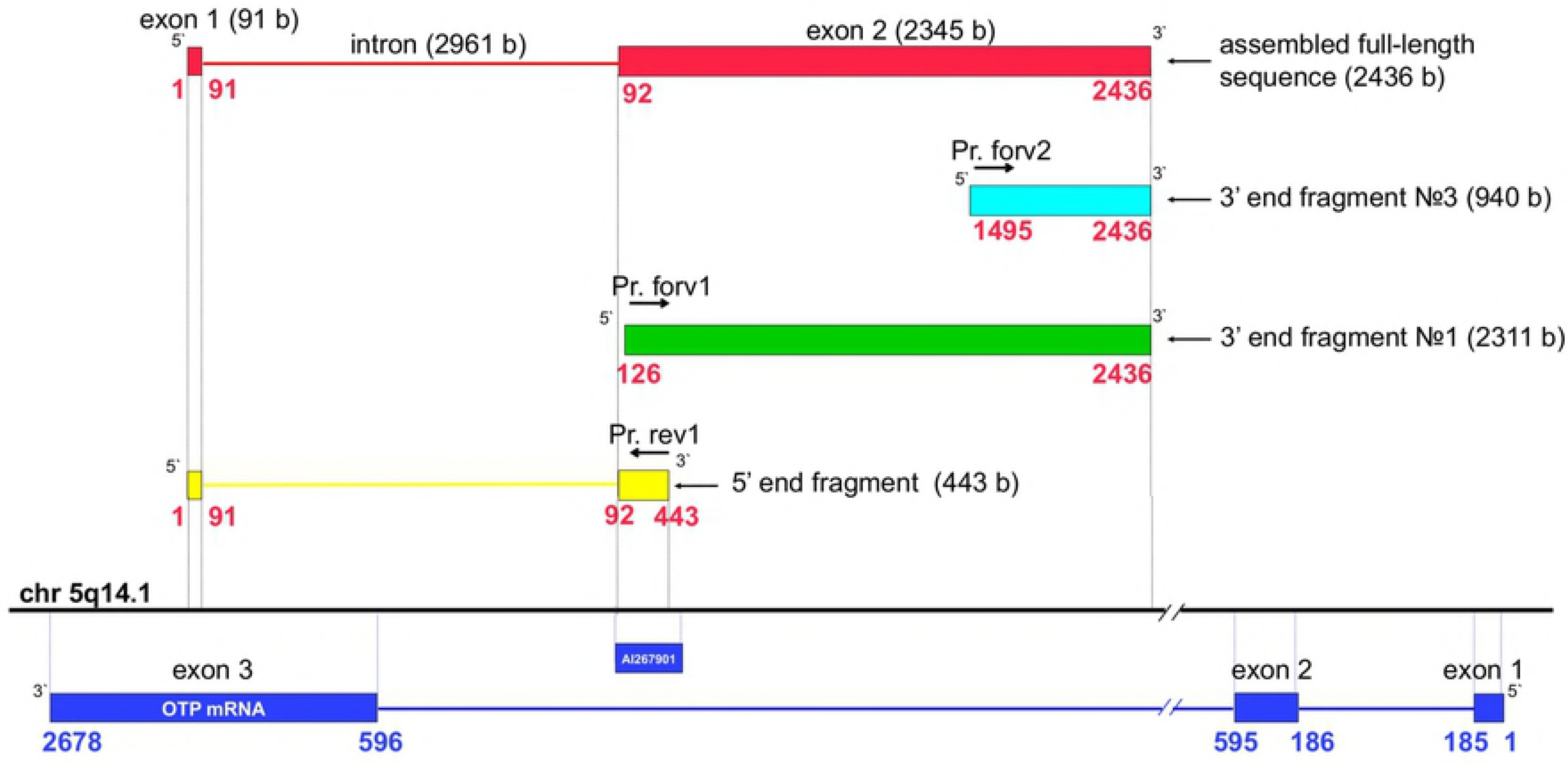
Two-round amplification of the full transcript on the cDNA from 1 - 293T cells 2 - uterus endothelium adenocarcinoma 3 - negative control, first round of PCR, no template added 4 - negative control, second round of PCR, no template added M1 - GeneRuler™ 100 bp DNA ladder (Fermentas) M2 - GeneRuler™ 1 kb DNA ladder (Fermentas)

The search of the possible open reading frames (ORFs) was performed using the ORF Finder webtool. We identified 10 ORFs coding for peptides from 20 to 62 amino acid long (S5 List).

Amino acid sequences of the identified ORFs were compared to the known proteins using Blastp algorithm. Homologous proteins were not found in humans or in other organisms.

Using RNAfold Web Server we modeled the secondary structure of the newly discovered gene mRNA (S6 Fig). This structure has a low free energy level (−650.96 kcal/mol) suggesting that this mRNA structure is thermodynamically stable.

We demostrated that different parts of our gene have different evolutionary ages. Orthologs of sequences located between 1-91st nucleotide and 600^th^ to 2436^th^ nucleotides were found in all *Tetrapods*, but othologs of sequence located between 92^nd^ and 600^th^ nucleotides were found only in *Eutherian* species, as seen on the phylogenetic trees (S7 and S8 Figs). The sequence conservation analysis was performed with Phylip tool integrated in USCS genome browser. We found that the sequence between 92nd and 600th nucleotides demonstrated low conservation in all genomes used for the analysis (Fig 6).

**Fig 6.**
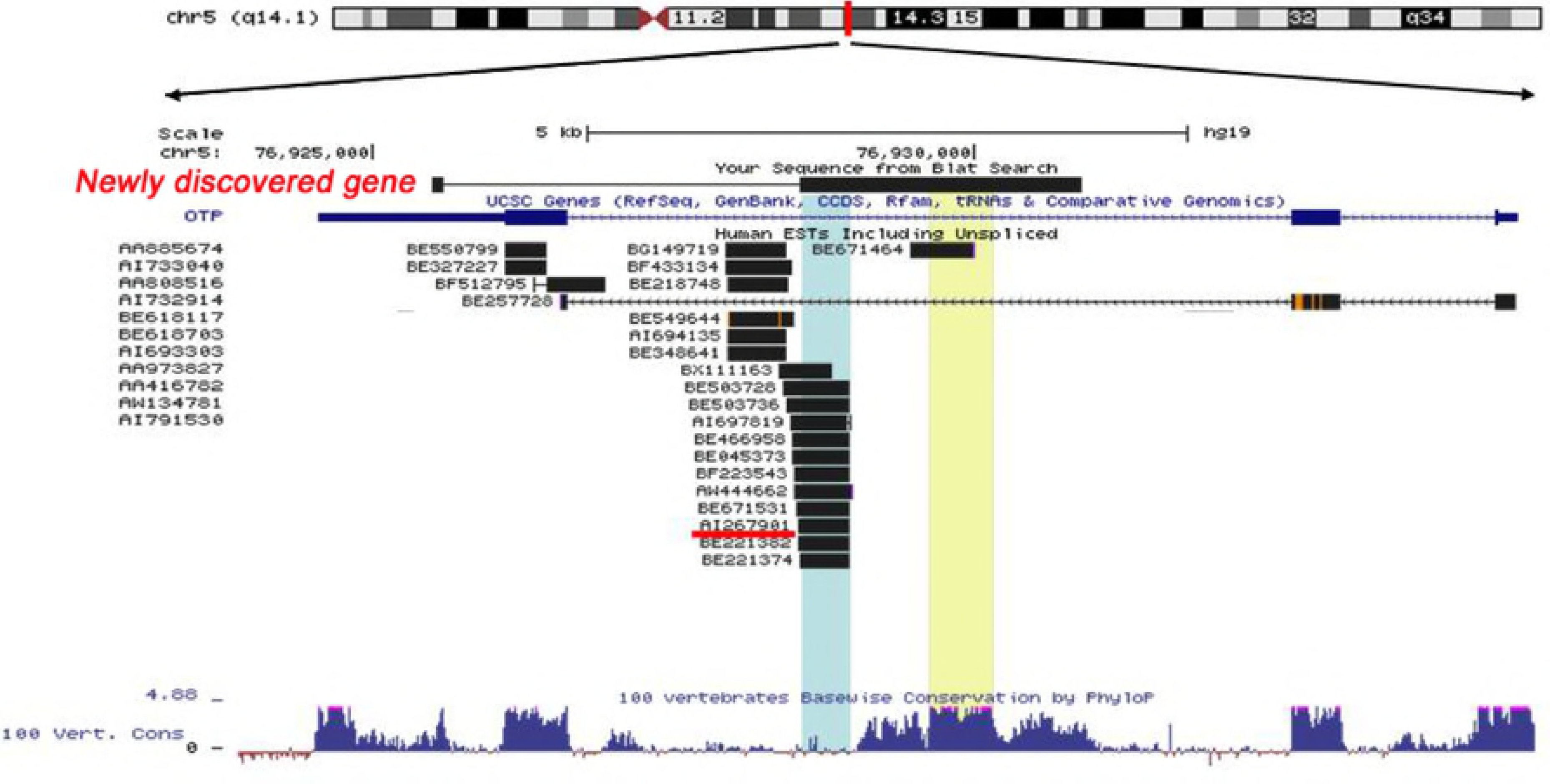
Chromosome location of *OTP* and newly discovered gene with mapping of the known ESTs. The least conserved genomic DNA sequence is highlighted with blue. The most conserved genomic DNA sequence is highlighted with yellow. (This scheme was obtained with UCSC Genome Browser tool).

To assess the tumor specificity of expression of the newly identified gene we used commercial Clontech (USA) and BioChain (USA) cDNA panels corresponding to normal and tumor tissues. We also used cDNA panel made in our laboratory using clinical samples of tumors from various localizations and at different stages of progression (Biomedical Center human tumor cDNA panel, Russia). The gene expression was determined by PCR with primers specific to the most conservative part of the newly discovered gene - from 1012 to 1452 nucleotide (Fig 6, in yellow). In normal and fetal tissue cDNA panels, the minor specific signal corresponding to this fragment was detected in only one testis sample (Fig 7).

**Fig 7.**
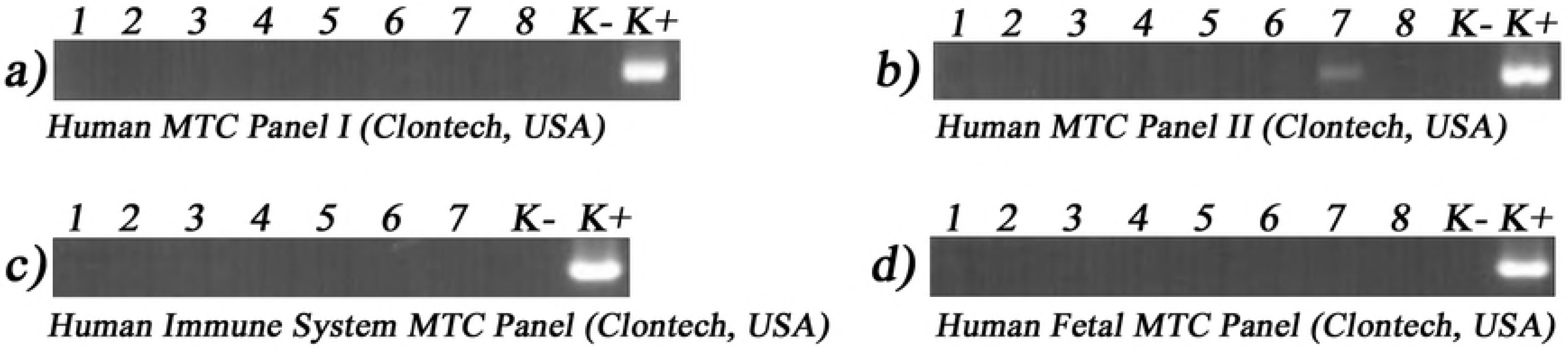
The lack of expression of the newly discovered gene conservative region in normal human tissues. a) 1 − normal brain, 2 − normal heart, 3 − normal kidney, 4 − normal liver, 5 − normal lung, 6 − normal pancreas, 7 − normal placenta, 8 − normal skeletal muscle, K- − PCR with no template, K+ - PCR with human DNA. b) 1 − normal colon, 2 − normal ovary, 3 − normal peripheral blood leukocytes, 4 − normal prostate, 5 − normal small intestine, 6 − normal spleen, 7 − normal testis, 8 − normal thymus, K+ − PCR with no template, K+ - PCR with human DNA. c) 1 − normal bone marrow, 2 − fetal liver, 3 − normal lymph node, 4 − normal peripheral blood leukocyte, 5 − normal spleen, 6 − normal thymus, 7 − normal tonsil, K- − PCR with no template, K+ − PCR with human DNA. d) 1 − fetal brain, 2 − fetal heart, 3 − fetal kidney, 4 − fetal liver, 5 − fetal lung, 6 − fetal skeletal muscle, 7 − fetal spleen, 8 − fetal thymus, K- − PCR with no template, K+ − PCR with human DNA.

In BioChain tumor cDNA panels, the expression of newly discovered gene was detected in the following tumors: carcinomas of lung and bladder, adenocarcinoma of esophagus, small intestine, colon and ovary. The fragment was not found in brain astrocytoma, testis seminoma, carcinomas of breast, liver, kidney, fallopian tube and ureter, stomach and uterus adenocarcinoma (Fig 8a).

**Fig 8.**
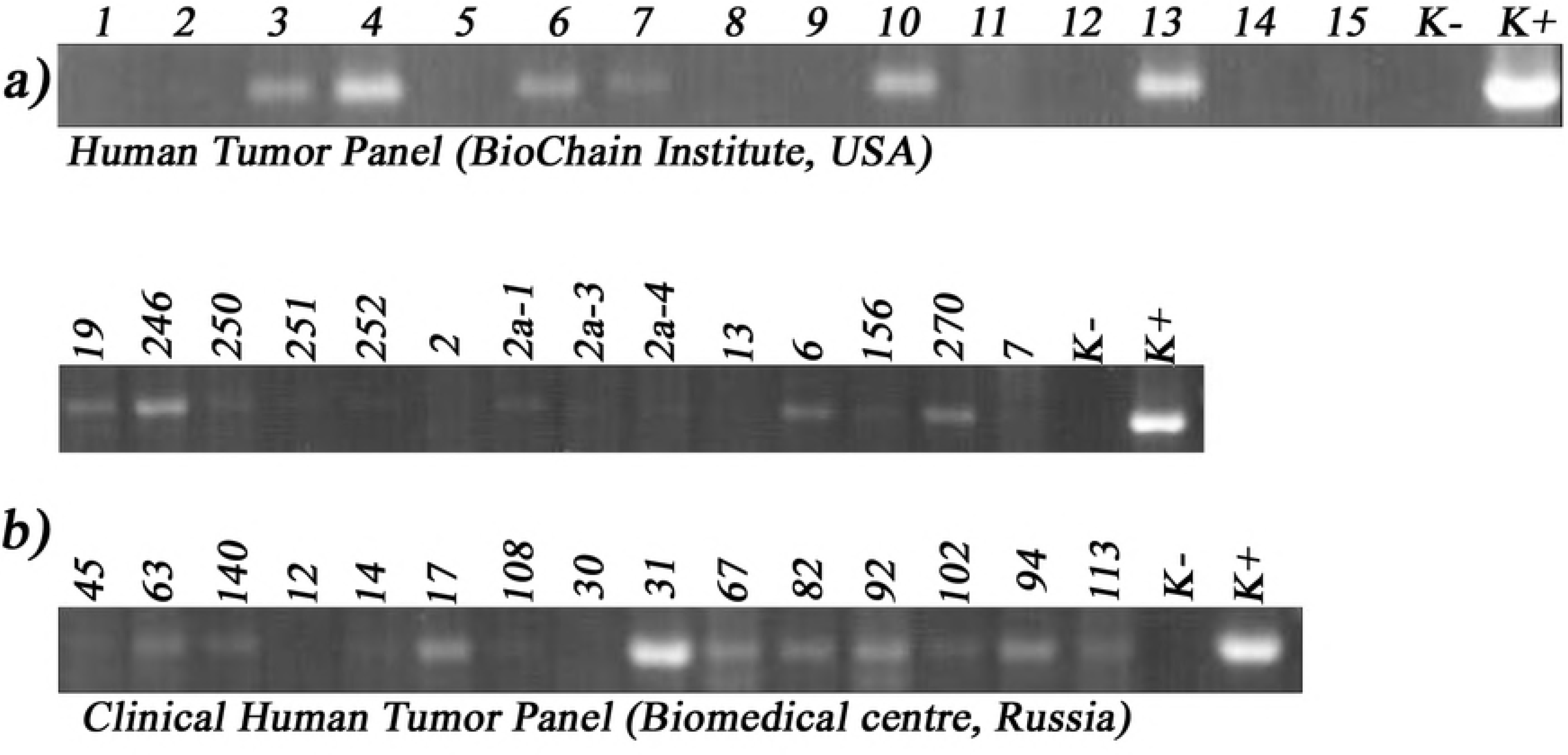
Expression of the newly discovered gene conservative region in human tumors. a) 1 - brain astrocytoma, 2 - breast invasive ductal carcinoma, 3 - lung squamous cell carcinoma,
4 - esophagus adenocarcinoma, 5 - stomach adenocarcinoma, 6 - small intestine adenocarcinoma, 7 - colon adenocarcinoma, 8- hepatocellular carcinoma, 9- kidney clear cell carcinoma, 10 - bladder transitional cell carcinoma, 11- uterus adenocarcinoma, 12 - fallopian tube medullary carcinoma, 13 - ovary mucinous adenocarcinoma, 14 − testis seminoma, 15- ureter papillary transitional cell carcinoma, K- − PCR with no template, K+ − PCR with human DNA. b) 19 - stage III mammary gland adenocarcinoma, 246, 250, 251, 252 - stage II-III invasive duct mammary gland cancer; 2 - squamous cell cervical carcinoma IV stage and its metastases into uterus (2a-1), left (2a-3) and right ovary (2a-4), 13- cervical myosarcoma, stage II-III, 6 - ovary
cancer, 156 - stage II moderately differentiated endometrial adenocarcinoma, 270 - stage III moderately differentiated endometrial adenocarcinoma with metastases, 7 - seminoma, 45, 63 - meningiomas, 140 -hypophyseal adenoma, 12,14 - squamous cell lung cancer, 17 - bronchus cancer III stage, 108 −stomach cancer, 30 - stage IV chronic lymphacytic leukemia, 31 - stage IV non-Hodgkin T-cell lymphoma, 67 - lymphoadenpathy of unclear pathogenesis, 82 - stage II non-Hodgkin lymphoma, stage II, 92 - stage IV Hodgkin’s lymphoma, 94 - hemolythic anaemia of unclear pathogenesis, 102 - stage II non-Hodgkin lymphoma, 113T - stage IV non-Hodgkin lymphoma, K- − PCR with no template, K+ − PCR with human DNA.

In the Biomedical Center human tumor cDNA panel the expression of the studied fragment was detected in five samples of mammary gland cancer (19, 246, 250, 251, 252); in one sample of ovary cancer (6), hypophyseal adenoma (140) and bronchus cancer (17); in two samples of endometrial adenocarcinoma (156, 270), meningiomas (45, 63); and in all lymphoma samples (31, 67, 82, 92, 102, 94, 113) (Fig 8b). Weak signals corresponding to the studied fragment were also found in samples of metastasis of squamous cell cervical carcinoma from uterus (2a-1), left (2a-3) and right ovaries (2a-4), and in one of two samples of squamous cell lung cancer (14). No signals were found in cervical carcinoma (2), cervical myosarcoma (13), seminoma (7), stomach cancer (108), leukemia (30), and in one of two samples of squamous cell lung cancer (12) (Fig 8b).

We also studied expression of the extended transcript of the newly identified gene on RNA isolated from different tumors (non-Hodgkin’s lymphoma at stages II and IV, lymphadenopathy of unknown origin, invasive ductal breast cancer at stage II). cDNA was synthesized using oligo(dT) primer. We conducted 2-round PCR with the primers to the ends of the newly identified transcript (as-forv, as-rev). Results are presented at Fig 9. The fragment of 2378 bp long was found in all studied samples, thus demonstrating the expression of the previously unknown gene in different tumors.

**Fig 9.**
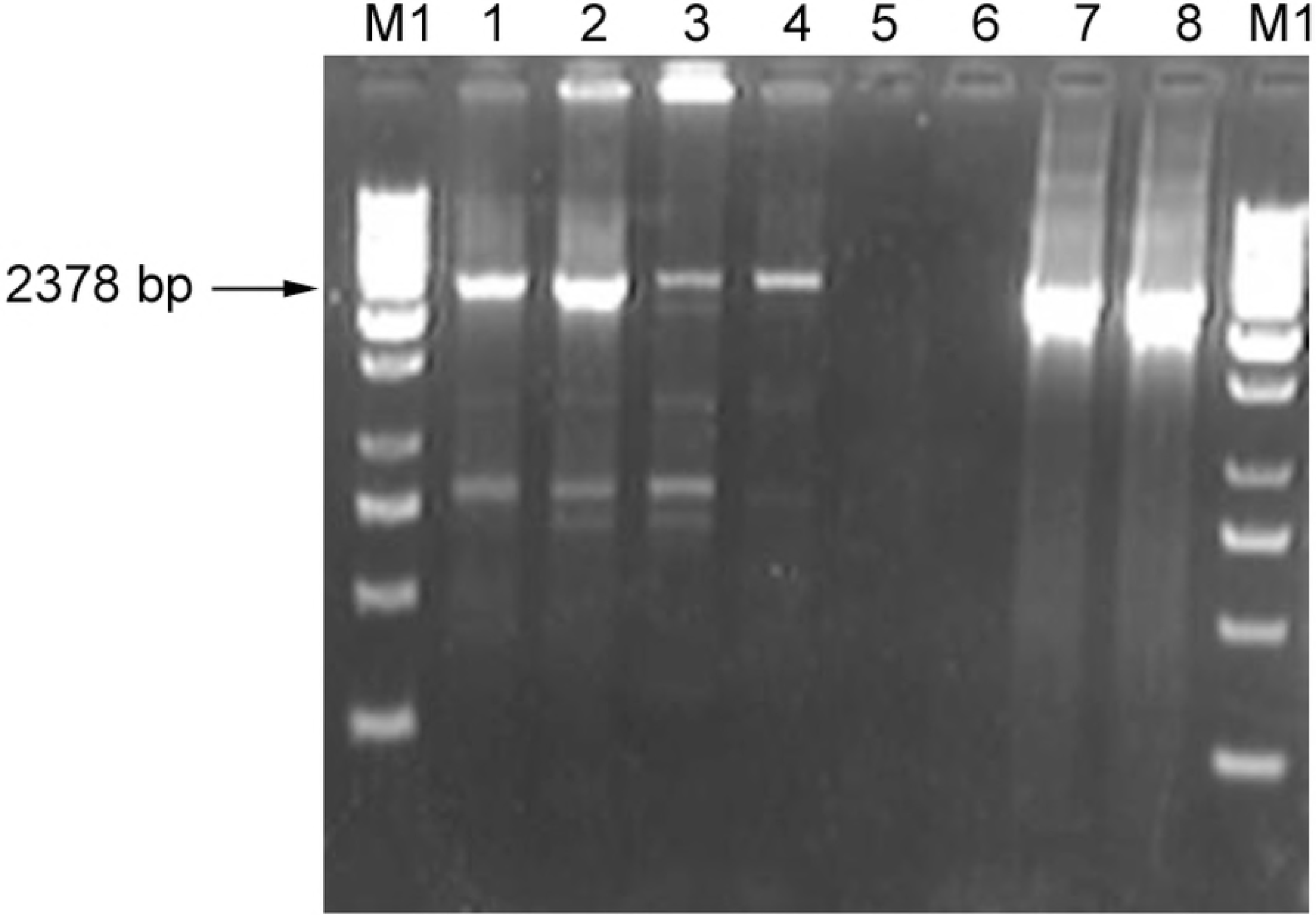
Two-round amplification of the studied transcript on the cDNA from differend clinical tumor subjects. 1- lymphadenopathy of unknown origine (67) 2- non-Hodgkin’s lymphoma at stage II (82) 3 non-Hodgkin’s lymphoma at stage IV (113) 4 - invasive ductal breast cancer at stage II (246) 6 - negative control, first round of PCR, no template added 7 - negative control, second round of PCR, no template added 8 − positive control, first round of PCR with plasmid containing full transcript of newly identified gene 9 − positive control, second round of PCR with plasmid containing full transcript of newly identified gene M1 - GeneRuler™ 1 kb DNA ladder (Fermentas)

## Discussion

In our previous papers [2, 3] we demonstrated highly tumor-specific expression of AI267901-like EST (earlier corresponding to cluster Hs.202247). This locus was expressed in 49 of 59 tumor samples of different localization and only in one sample of normal testis. We mapped the AI267901 sequence to the human genome using UCSC Genome Browser. The studied sequence was found to be located at 5q14.1 in the second intron of the human *Orthopedia homeobox (OTP)* gene (Figs 4 and 6).

Human *OTP* gene is the homologue of the murine *Orthopedia* gene and belongs to the homeodomain genes. Its function in humans is not fully characterized as yet. Expression of the *OTP* gene was found in the brain of 17 week old human embryo, supposing the potential role of this gene in brain development [4]. *OTP* is encoded by the “minus” strand of the 5th chromosome and consists of 3 exons and 2 introns.

At the present time *OTP* gene is actively studied as a prognostic marker for carcinoid lung tumors by several research groups. There are data showing the *OTP* gene expression only in lung tumors [5, 6] and bladder [6]. We have shown that *OTP* gene is expressed in a wide range of tumors of almost all localizations. This gene was expressed in 23 of 29 tumor samples of different localization and only in one sample of normal testis [3].

AI267901 is located in the intronic region of the *OTP* gene and must be absent in the mature *Orthopedia homeobox* mRNA. At the same time, our results show that AI267901 and *OTP* gene have similar expression profiles both in normal tissues and in tumors [3]. Therefore the AI267901 might be interpreted either as *OTP* transcript alternatively processed in tumors or as a separate transcript.

To find the complete nucleotide sequence of AI267901 we used the Rapid Amplification of cDNA Ends (RACE) and other approaches. As a result, we have obtained a 2436 bp long sequence of the previously unknown gene located on the opposite strand of *OTP* gene. mRNA of this gene is polyadenylated and has two exons. The first short exon (91 bp) is mapped to the antisense strand of untranslated region (3’-UTR) of *OTP* gene. It is known that initiation sites of noncoding RNAs transcription are frequently located in the 3’-UTRs of protein-coding genes [7]. The second exon (2345 bp) is mapped to the antisense strand of 2^nd^ intron of *OTP* gene. The first and second exons are separated by 2961 bp intron (Figs 4 and 6). The sequence of the whole transcript is presented in S4 Sequence.

We have shown the expression of the full size newly discovered gene in human embryonic kidney *293* (HEK293) cell line and in uterus endothelium adenocarcinoma (Fig 5), and of the 2378 bp fragment of the gene in the following human tumors: non-Hodgkin lymphoma stage II and stage IV, invasive ductal breast cancer stage II and lymphoadenpathy of unknown origin (Fig 9). Interestingly that several shorter sequences can be seen along with the expected fragment (2378 bp) at Fig 9. These fragments might be an alternatively spliced variants of the newly discovered gene in tumors. These fragments are being sequenced now in our laboratory.

The study of the expression of the conservative region of newly discovered gene in human tumors and normal tissues has shown that it was expressed in majority of the tumor samples studied, including tumors of brain, lung, esophagus, intestines, breast, bladder, uterus, ovary and in lymphomas (Fig 8), but among normal tissues only in testis (Fig 7).

The search of possible open reading frames (ORFs) was performed using the ORF Finder webtool. Amino acid sequences of the ORFs were short in comparison with the known proteins. Blastp algorithm did not show homologous proteins in humans or in other organisms.

The lack of significant ORFs suggests that we discovered a new tumor-specific long noncoding RNA (lncRNA). Thus, *OTP*- antisense RNA 1 gene is supposed to be a cancer-testis (CT) long non-coding RNA (lncRNA) gene. According to human non-protein coding RNA (ncRNA) gene nomenclature [8, 9] we assigned a symbol *OTP-AS1* (*OTP* − antisense RNA 1) to this gene. This gene symbol is approved by HUGO Gene Nomenclature Committee (HGNC).

We found that, despite the fact that most of the gene (~1900 nucleotides) appeared in *Tetrapoda*, the insertion of evolutionarily novel part occurred after *Eutheria* speciation (S8 Fig). This evolutionarily novel part locates between 92^nd^ and 600^th^ nucleotides on *OTP-AS1* sequence

Most lncRNAs exhibit weak or untraceable primary sequence conservation [10, 11, 12]. Nevertheless conservative lncRNAs are described in the literature, e.g. *MALAT1* gene [13]. *OTP-AS1* older part is also conservative as shown by Phylop analysis (Fig 6). The evolutionarily younger part (~500 nucleotides) of the *OTP-AS1* gene demonstrates much lower level of conservation according to analysis made by Phylop tool (Fig 6). Interestingly, all AI267901-like tumor-specific ESTs are mapped on this non-conserved *Eutheria*-specific region of the gene. This links *OTP-AS1* gene to so-called *TSEEN* (tumor-specifically expressed, evolutionary novel) genes described by the authors in previous papers [14, 15, 16, reviewed in 17].

The *OTP-AS1* gene transcript is spliced and polyadenylated. mRNA of *OTP-AS1* gene shows low levels of free energy (−650.96 kcal/mol). This means that thermodynamic stability of the studied RNA is high. This possibly indicates to the existence of some functions of this gene.

Although functions of most lncRNAs are unknown, the number of characterized lncRNAs is growing and many publications suggest they play roles in regulation of gene expression in development, differentiation and human disease. lncRNAs may regulate protein-coding gene expression on both transcriptional and posttranscriptional levels [reviewed in 18].

Cancer-testis (CT) genes demonstrate similarity between processes of spermatogenesis and tumorigenesis. CT-antigens may serve as cancer diagnostic markers or as potential targets for anti-cancer vaccines. The large group of cancer-testis non-coding RNA (CT-ncRNA) was recently described by *in silico* methods [19]. Wang and co-authors described cancer-specific CT-coding gene/CT-ncRNA pairs (where the distance between the CT-coding gene and CT-ncRNA was <100 kb). The authors suggest that these pairs may be involved in self regulatory interactions. For example, it was demonstrated that meiosis-related extremely highly expressed CT genes (MEIOB) and their companion testis-specific ncRNAs (TS-ncRNA; LINC00254) play crucial roles in carcinogenesis in lung adenocarcinoma [19].

According to our findings in this paper and elsewhere [3], *OTP-AS1* gene as well as *OTP* gene may be CT-genes. Moreover, *OTP-AS1* gene is located at antisense to *OTP* gene strand. Thus, *OTP-AS1* and *OTP* genes may be the CT-coding gene/CT-ncRNA pair involved in regulatory interactions. This is supported by the similar profile of their expression [3]. These genes may be targets for future cancer therapies.

Thus we have discovered a new cancer-testis long noncoding RNA which may have regulatory function. We assume that *OTP-AS1* gene may play a role in *OTP* gene regulation.

Part of this data was presented at the 2nd International Conference on the Long and the Short of Non-Coding RNAs (09.06.2017 - 14.06.2017, Heraklion, Crete, Greece)

## Materials and Methods

### cDNA Panels

#### MTC™ Panels

For studies of gene expression we used commercial cDNA panels. The panels containing a set of normalized single-strand cDNA, produced from poly(A)+ RNA from various normal human tissues were obtained from Clontech, USA. We used the following panels: Human MTC™ Panel I (Catalog no. 636742), Human MTC™ Panel 2 (Catalog no. 637643), Human Immune System MTC™ Panel (Catalog no. 636748) end Human Fetal MTC™ Panel (Catalog no. 636747). According to the manufacturer’s information, the panels were free from genomic DNA and were normalized to expression levels of four house-keeping genes. Each cDNA sample comes from a pool of tissue samples obtained from donors of different age and sex, with 2-550 donors in each pool, and the fetal tissue samples were obtained from spontaneously aborted fetuses at 18 to 36 weeks of gestational age. The quality of all samples we assessed by PCR using primers for the housekeeping gene *GAPDH* (data not shown).

### Tumor cDNA Panel

A cDNA panel containing a total of 15 of cDNA samples were obtained from BioChain Instutute, USA (Catalog nos.: C8235544, C8235545, C82355546, C8235549). The samples were produced by the manufacturer from various human tumors obtained from surgeries. Each sample came from one patient and was histologically characterized. cDNA was produced from poly(A)+ mRNA that was free from genomic DNA and normalized by β-actin gene expression level. The quality of all samples we assessed by PCR using primers for the housekeeping gene *GAPDH.*

### Clinical Material

In our work we also used samples of surgically excised tumors of various origins obtained in the Kirov Military Medical Academy (St. Petersburg, Russia). A total amount of 29 samples was obtained in our laboratory after a written informed consent of all participant patients. The use of the samples for gene expression studies was approved by the Ethical Committee of the Kirov Military Medical Academy and the Biomedical Centre (St. Petersburg, Russia). The tumors were histologically characterized. We studied the following tumor samples: stage II-III invasive duct mammary gland cancer (3 samples, patient codes: 250, 251, 252), stage III mammary gland adenocarcinoma (patient code 19), squamous cell cervical carcinoma, IV stage (patient code 2) and its metastases into uterus (patient code 2a-1), left (patient code 2a-3) and right ovary (patient code 2a-4), cervical myosarcoma, stage II-III (patient code 13), ovary cancer (patient code 6), moderately differentiated endometrial adenocarcinoma, stage II (patient code 156), moderately differentiated endometrial adenocarcinoma with metastases, stage III (patient code 270), seminoma (patient code 7), meningioma (patient codes 45, 63), hypophyseal adenoma (patient code 140), squamous cell lung cancer (patient codes 12, 14), bronchus cancer III stage (patient code 17), stomach cancer (patient code 108), chronic lymphacytic leukemia, stage IV (patient code 30), non-Hodgkin T-cell lymphoma, stage IV (patient code 31), lymphoadenpathy of unclear pathogenesis (patient code 67), non-Hodgkin lymphoma, stage II (patient code 82), Hodgkin’s lymphoma, relapse, stage IV (patient code 92), hemolythic anaemia of unclear pathogenesis (patient code 94), non-Hodgkin lymphoma, stage II (patient code 102), non-Hodgkin lymphoma, stage IV (patient code 113), invasive ductal breast cancer at stage II (patient code 246). Additionally, we used human embryonic kidney 293 (HEK293) cell line.

### RNA isolation and quality control

Total RNA isolation from clinical material of human tumors was performed via routine procedure using guanidine isothiocyanate as described elsewhere [20]. RNA samples were treated with DNAse I, RNAse free (Sigma, USA) for 10 min at 25°C in order to remove any contaminating genomic DNA.

The concentration of isolated RNA was measured using Ultrospec^®^ 3100 *pro* spectrophotometer. RNA quality was assessed spectrally by the A260/A280 ratio and visually following agarose gel electrophoresis by band intensity ratio of 28s rRNA to 18s rRNA [21].

The absence of the DNA in the RNA samples was determined by PCR using primers for the housekeeping gene *GAPDH* (forward 5’ - TGAAGGTCGGAGTCAACGGATTTGGT -3’, reverse 5-CATGTGGGCCATGAGGTCCACCAC -3). Conditions for PCR amplification were as follows: 3 min of denaturation at 94 °C; 40 cycles of 30 s at 94 °C, 30 s at 68 °C, 30 s at 72 °C; followed by a final extension for 5 min at 72 °C. The resulting amplificates were resolved by electrophoresis in 2% agarose gel and stained with ethidium bromide. The absence of the DNA contamination in RNA samples was indicated by the lack of the 983 bp long amplification product of *GAPDH*. The gels were photographed under UV illumination.

### RACE (Rapid Amplification of cDNA Ends)

We used MarathonTM cDNA Amplification Kit (Clontech) to obtain cDNA from uterus adenocarcinoma RNA samples according to manufacturer’s protocol. Obtained two-strand cDNA was subjected to 5’- and 3’- RACE PCR.

5’ - RACE PCR is two-round amplification using the gene specific forv1 (5’- CGATGGATAAACAGGTCTCGTCTCTTCC-3’, T_m_=62°C), forv1N (5- AGGTCTCGTCTCTTCCCAGTTGCAG-3’ T_m_= 61°C) and adaptor (5’- GGCCAGGCGTCGACTAGTAC-3’) primers.

3’ - RACE PCR is two-round amplification using the gene specific rev1 (5’- TGCAGGTTGTTAGGAACCGGTCTTG-3’ T_m_=62°C), rev1N (5-TTAGGAACCGGTCTTGATTTTATAAGAC -3’ T_m_=56°C) and adaptor primers.

Gene specific primers (forv1, rev1) were designed to provide the overlapping of the obtained PCR products.

The PCR mixture contained 2μl of 1:50 cDNA dilution, PCR-buffer (Qiagen, Germany), 100 μM (each) dATP, dGTP, dTTP and dCTP, 5 pmol of each primer, and 1 unit of Hot Taq DNA polymerase (Qiagen, Germany) in a total of 25-μl reaction volume.

Conditions of the reactions were as follows: 15 min of denaturation at 94 °C; 10 pre-cycles of 30 s at 94°C, 30 s at 68°C, 4 min at 72°C. Then 5 pmol of the adaptor primer was added to the reaction mix and amplification was continued under the following conditions: 1 min of denaturation at 94°C; 5 cycles of 30 s at 94°C and 4 min at 72°C, 5 cycles of 30 s at 94°C and 4 min at 70°C, 25 cycles of 30 s at 94°C and 4 min at 68°C; followed by a final extension for 5 min at 72 °C.

A 50-fold diluted 1 μl aliquot from the 1^st^ round of amplification was used for the 2^nd^ round of amplification with nested primers under the same conditions, but without the 10 precycles. The resulting amplificates were resolved by electrophoresis in 2% agarose gel and stained with ethidium bromide.

### Determination of the 3’ end of the transcript

To determine the 3’ end of the transcript reverse transcription (RT) followed by 2-round PCR were performed. cDNA from 293T cells RNA was obtained using SuperScript™III Reverse Transcriptase (Invitrogen) with oligo(dT) adapter primer (5’- GGCCAGGCGTCGACTAGTACTTTTTTTTTTTTTTTTT-3’) according to manufacturer’s protocol. The cDNA was subjected to 2-round amplification with adapter and gene-specific primers forv1 and forv2 (5’- GTGCAGAAGTTATTTTACTGATTTG -3’ T_m_=63°C). The PCR mixture contained 1μl of cDNA, PCR Taq buffer (Invitrogen), 3 mM MgCl_2_, 100 μM (each) dNTP, 5 pmol of each primer, and 1 unit of Platinum Taq DNA Polymerase (Invitrogen) in a total of 25-μl reaction. Conditions of the reactions were as follows: 2 min of denaturation at 94 °C; 5 cycles of 15 s at 94°C, 15 s at 60°C, 5 min at 72°C, 35 cycles of 15 s at 94°C, 15 s at 55°C, 5 min at 72°C; and a final extension for 5 min at 72 °C.

A 2 μl aliquot from the 1^st^ round was used for the 2^nd^ round of amplification with nested primers forv1N, forv2N (5’- GTTATTTTACTGATTTGGTTTTTATG -3’ T_m_=63°C) and adapter primer under the same conditions, but with Tm increased by 2 degrees (62°C and 57°C, respectively). The resulting amplificates were resolved by electrophoresis in 2% agarose gel and stained with ethidium bromide.

### cDNA synthesis

To obtain the full-length PCR-product of the studied gene we used cDNA prepared with SuperScript™ III Reverse Transcriptase (Invitrogen) with oligo(dT) primer on RNA from 293T cells and human tumors (patient codes 67, 82, 113, 156 and 246). cDNA was prepared as recommended by manufacturer.

To obtain the Biomedical Center human tumor cDNA panel we used Revert Aid^®^ First Strand cDNA Synthesis Kit (Fermentas, Lithuania) with random hexamer primers on RNAs from different human tumors, following the manufacturer guidelines.

Obtained cDNA samples were stored at −20°C. The quality of the samples was assessed by PCR using primers for the housekeeping gene *GAPDH* (data not shown).

### Two-round amplification of the full-length transcript

Two-round PCR was performed with the primers for the obtained 5’end and 3’end: as-forv (5’- TGCACAGCATGCCCTAGAC-3’ T_m_=60°C), as-rev (5’- TTTTACTGATTTGGTCATTATG-3’ T_m_=61°C) and full-forv (5’- GTCTGAGCGTGAGCGAGAG-3’ T_m_=62°C), full-rev (5’- ATGAAAAAAGAAAACGAGGTCTATT -3’ T_m_=58°C).

The PCR mixture contained 1μl of cDNA, PCR-buffer High Fidelity (Invitrogen), 2mM MgSO_4_, 100 μM (each) dNTP, 5 pmol of each primer, and 1 unit of Platinum DNA Polymerase High Fidelity (Invitrogen) in a total of 25-μl reaction. The first round of amplification was performed under the following conditions: 2 min of denaturation at 94 °C; 40 cycles of 15 s at 94°C, 20 s at 55°C, 3.5 min at 68°C; and a final extension for 5 min at 68 °C. A 1 pl aliquot from the 1^st^ round was used for the 2^nd^ round of amplification under the same conditions. The resulting amplificates were resolved by electrophoresis in 2% agarose gel and stained with ethidium bromide.

### PCR

PCR primers targeting *OTP-AS1* conservation region were designed based on our sequence of *OTP-AS1* from S1 Sequence. Forward primer 1012forv: 5’- CACTTTCATGATATCTGCTGTTAC-3’, reverse primer 1452rev: 5’- ATAGTGTGCTGTAATTCCATTG-3’. The expected size of the amplicon was 440 bp (from 1012 nucleotide to 1452 nucleotide of *OTP-AS1* sequence).

The PCR mixture contained 2.5 μl of cDNA, PCR-buffer (67 mM Tris-HCl, pH 8.9, 4 mM MgCl_2_, 16 mM (NH_4_)SO_4_, 10 mM 2-mercaptoetanol), 200 μM (each) dNTP, 1 unit of Taq DNA polymerase (Fermentas, Lithuania), and 5 pmol of each primer in a total of 25-μl reaction volume. Amplification was performed with the following conditions: 1 min at 95°C; 35 cycles consisting of 30 s at 95°C, 30 s at 60°C, and 60 s at 72°C; and final elongation at 72°C for 5 min.

All PCR products were analyzed by electrophoresis in 2% agarose gel and detected by staining with ethidium bromide.

### Sequencing

PCR-products were extracted from the agarose gel, cloned into the pGEM-T Easy Vector (Promega) or TA cloning vector (Invitrogen), propagated in E.coli and sequenced using conventional techniques.

### Software and databases

We used BioEditsoftware for basic manipulations with nucleic and amino acids sequences. Resources of the NCBI databases (http://www.ncbi.nlm.nih.gov/) and UCSC Genome Browser (GB) (http://genome.ucsc.edu/) were used extensively.

ORF search was performed with ORF Finder webtool (http://www.bioinformatics.org/sms2/orf_find.html). To analise the evolutionary age of the studied gene we used the bidirectional best hits (BBH) method, which entails identifying the pairs of genes in two different genomes that are more similar (i.e., with highest alignment score) to each other than to any other gene. The orthologs were searched in 40 completely sequenced eukaryotic genomes which were retrieved from “Genome” resource (http://www.ncbi.nlm.nih.gov/genome/). The list of the used genomes is in S9 List. The nHMMER tool [22] and the original Shell script were used to perform homology search. For nHMMER an e-value threshold of *1e*^−*10*^ was specified. The first hit for query sequence within the program output was considered as the best hit. The very same procedure was performed for the results ran in the opposite direction, i.e. for the results where the subject genome was used as a query, and the query genome was used as a subject. Phylogram and cladogram of complete transcript and of its Eutheria-specific part were obtained with MrBayes (v3.2.3) tool [23, 24]. More than 1000 iterations were implemented for bootstrap procedure. Phylop tool integrated in USCS genome browser was used for conservation analysis.

The secondary structure of gene mRNA was modeled with RNAfold Web Server (http://rna.tbi.univie.ac.at/cgi-bin/RNAWebSuite/RNAfold.cgi) [25, 26].

## Supporting Information

**S1 Sequence. Sequence of the 5’ end of the transcript.**

**S2 Sequence. Sequence of the fragment №1 of the 3’ end of the transcript.**

**S3 Sequence. Sequence of the fragment №3 of the 3’ end of the transcript.**

**S4 Sequence. Sequence of the the full lenght transcript of the study gene (*OTP-AS1*).**

**S5 List. Amino acid sequences of ORFs.**

**S6 Figure. Secondary structure of *OTP-AS1* mRNA.**

**S7 Figure. Phylogram of complete sequence of *OTP-AS1* gene orthologs based on average branch lengths.**

**S8 Figure. Phylogram of the evolutionary novel part of *OTP-AS1* gene orthologs based on average branch lengths.**

